# Structural insights into IMP2 dimerization and RNA binding

**DOI:** 10.1101/2024.02.16.580656

**Authors:** Stephen Zorc, Paola Munoz-Tello, Timothy O’Leary, Xiaoyu Yu, Mithun Nag Karadi Giridhar, Althea Hansel-Harris, Stefano Forli, Patrick R. Griffin, Douglas J. Kojetin, Raktim N. Roy, Michalina Janiszewska

## Abstract

IGF2BP2 (IMP2) is an RNA-binding protein that contributes to cancer tumorigenesis and metabolic disorders. Structural studies focused on individual IMP2 domains have provided important mechanistic insights into IMP2 function; however, structural information on full-length IMP2 is lacking but necessary to understand how to target IMP2 activity in drug discovery. In this study, we investigated the behavior of full-length IMP2 and the influence of RNA binding using biophysical and structural methods including mass photometry, hydrogen-deuterium exchange coupled to mass spectrometry (HDX-MS), and small angle x-ray scattering (SAXS). We found that full-length IMP2 forms multiple oligomeric states but predominantly adopts a dimeric conformation. Molecular models derived from SAXS data suggest the dimer is formed in a head-to-tail orientation by the KH34 and RRM1 domains. Upon RNA binding, IMP2 forms a pseudo-symmetric dimer different from its apo/RNA-free state, with the KH12 domains of each IMP2 molecule forming the dimer interface. We also found that the formation of IMP2 oligomeric species, which includes dimers and higher-order oligomers, is sensitive to ionic strength and RNA binding. Our findings provide the first insight into the structural properties of full-length IMP2, which may lead to novel opportunities for disrupting its function with more effective IMP2 inhibitors.

## INTRODUCTION

RNA Binding Proteins (RBPs) play multiple critical roles in normal human physiology as well as disease, as they regulate key cellular processes by controlling RNA localization, stability, translation, and splicing (1–3). Their involvement in cell proliferation, motility and metabolism is necessary for homeostasis, while dysfunctional RBPs can contribute to a wide array of pathologies, including diabetes, cancer, cardiovascular disease, and neurodegeneration (1–3).

Insulin-like growth factor 2 mRNA binding protein 2 (IMP2, IGF2BP2) is an RNA binding protein that belongs to the IMP family with two other paralogs, IMP1 and IMP3 (4, 5). It is involved in RNA stability, localization, and translation, and has been implicated in protection of pluripotency-associated transcripts from miRNA-mediated degradation (4). IMP2 overexpression has been linked to a wide array of malignancies, including glioblastoma, melanoma, and breast cancer, where it supports the survival of cancer stem-like cells and cancer cell metastasis by controlling stability and transport of distinct mRNAs (6–9). Moreover, IMP2 has also been linked to type 2 diabetes, as its control of mRNA stability is essential for maintenance of adipose tissue homeostasis (10, 11).

IMP2 consists of modular domain architecture containing six RNA-binding domains (RBDs), two RNA recognition motifs (RRMs), and two pairs of hnRNP K homology domains (KH1-2, KH3-4). IMP2 shares 56% of its sequence with its paralogs IMP1 and IMP3, yet it regulates a distinct pool of RNAs (12). Structures of individual IMP2 domains have been determined for RRM1 and KH3-4 domains; however, structural studies on full-length IMP2 have not been reported. Suzuki et al. obtained an NMR structure of RRM1 (residues 2-81, PDB: 2CQH), which covers about 13% of the entire protein sequence. Biswas and colleagues published an X-ray structure of KH34 domains (residues 426-588; PDB: 6ROL) which encompasses 27% of the full-length protein (12). AlphaFold structure prediction of full-length IMP2 shows that much of the protein is disordered. Besides the structured domains (RRM1, RRM2, KH1, KH2, KH3, and KH4), there are long stretches of unstructured regions connecting the RRM and KH domains that display low (50–70) or very low (< 50) per-residue AlphaFold prediction confidence scores.

Regardless of organism or cell type, all members of the IGF2BP protein family have been shown to bind RNA (4, 5). *In vitro* studies have demonstrated that RNA-binding is facilitated via the KH-domains in IMP1, although the RRM domains could potentially contribute to the stabilization of IGF2BP-RNA complexes, increasing their *in vitro* half-life (13). Recent structural analyses of human IGF2BP1 KH3/4 suggest the formation of an anti-parallel pseudo-dimer conformation of the KH3 and KH4 domains, each contacting a strand of RNA (14). However, there are no studies that describe the oligomeric states of full-length IMP2 and how RNA binding influences full-length IMP2 structure.

In recent years, IMP2 has gained attention as a promising molecular target in various cancers (8, 15, 16). However, the moderate potency of the IMP2 targeting small molecules optimized based on partial domain information points to the importance of further understanding of the structure and function of full-length IMP2 (17, 18). In this study, we found that the primary oligomeric state of full-length IMP2 is a homodimer and determined regions of IMP2 involved in RNA interaction. Using a variety of biophysical and structural methods including mass photometry, hydrogen-deuterium exchange coupled to mass spectrometry (HDX-MS), and small angle X-ray scattering (SAXS). Our findings provide a refined understanding of the solution behavior of IMP2 in presence and absence of RNA. Furthermore, we describe the equilibrium of the various states of IMP2, and how it can be shifted by ionic strength and RNA binding.

## RESULTS

### IMP2 adopts a dimeric form in solution

During the protein purification of full-length IMP2, expected to be ^∼^66kDa, we noticed multiple peaks in size exclusion chromatography (SEC) profiles (**Fig. S1**), suggesting that IMP2 exists in multiple oligomeric states. Of particular interest was the peak corresponding to the size of an IMP2 dimer, which was largely abundant. While IMP2 has been considered to be a monomeric protein and modeled as such in previous research studies, the prevalence of oligomeric species has been discussed for other IMP family members (13, 17, 19). To gain insight into the structural properties of full-length IMP2 - 3D molecular envelope, molecular mass, radius of gyration and other structural attributes - we collected in-line SEC-SAXS data. Molecular weight simulation from the SAXS data confirmed the presence of the dimeric IMP2 state (127 kDa). We also studied the oligomeric state of IMP2 using mass photometry and SEC-multi-angle light scattering (SEC-MALS) and observed the presence of multiple IMP2 species (*vide infra*), but most notably the dimeric form.

After data collection, the SAXS molecular envelope was fitted with a protein dimer model with a *χ*^2^ value of 3.46 (**Fig. 1A-C**). Alternate models with different domain locations displayed high *χ*^2^ values and were therefore ruled out as unlikely structures due to deviations from the theoretical scattering profiles. Our primary observation from in-line SEC-SAXS data indicates that IMP2 is present predominantly in a dimeric form in the absence of RNA, harboring an asymmetric configuration, where the KH12 domains from the first molecule are more separated from the rest of the complex than the equivalent KH12 domains from the second molecule (**Fig. 1B**). We designated each molecule with a superscript number: ^1^ would present domains from the first molecule and ^2^ denotes domains of the second protein molecule within the dimer. The interaction between KH12^2^ and KH34^2^ reveals a continuous buried surface. In the first molecule, the KH12 domain is separated from the KH34 domain by ∼ 50Å. This is evident from the SAXS envelope, with a signature protuberance at the center that fits the KH12 domain. Our fit for theoretical scattering curve, for this configuration of the domains also matched our experimental scattering results (with a *χ*^2^ fit of 3.46). Although our SAXS-determined structural model is devoid of the connecting loop regions, the extensively floppy attributes of these intrinsically disordered regions add minimal value to the uniqueness of the structural envelope.

**Figure 1.**
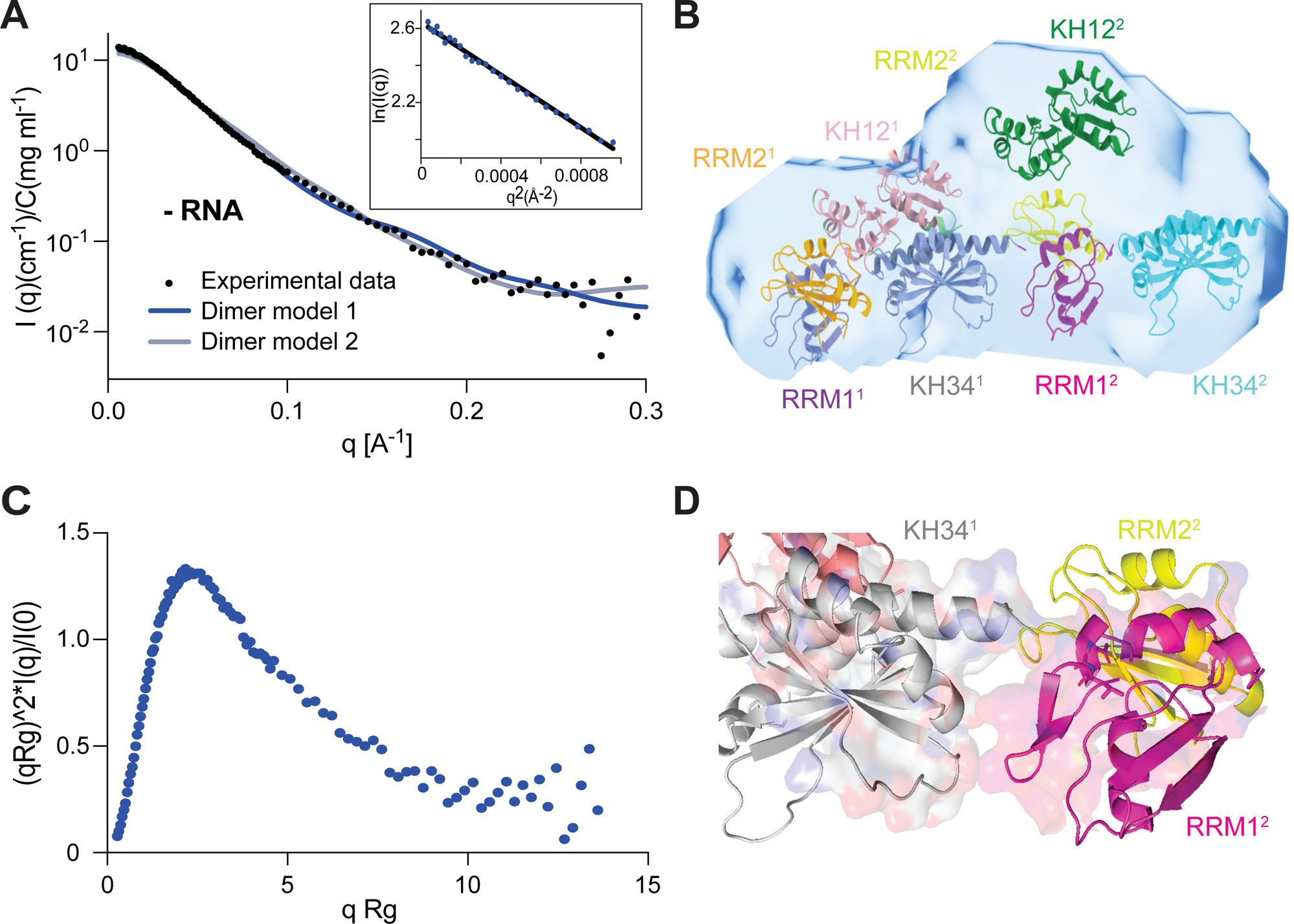
IMP2 primarily exists as a dimer in solution. **A)** The log I(q) versus q plot represents the primary SAXS data, with Guinier plots shown as inset. The maximum dimension is 142 Å, and the minimum q measured (0.006 Å^−1^) is well below the minimum recommended for accurate assessment. Importantly, in both experiments, the Guinier plots (inlay) are linear to the first measured q values and a plot of log I(q) versus log q shows that the slope is effectively zero at low q as expected for monodisperse particles of similar size. These measures together provide confidence that the data are free of significant amounts of contaminating species or inter-particle correlations contributing a structure-factor term to the scattering. **B)** SAXS envelope with apo/RNA-free dimer model within. **C)** Dimensionless Kratky plot demonstrates that the SAXS data are from predominantly folded particles. The IMP2 - RNA plots display the expected bell-shaped curves, with a maximum of about 1.3 at around qR_g_ = 2.07. The small rise of IMP2 apo/RNA-free evident at qR_g_ > 11 suggests some flexibility. **D)** Zoomed in portion of apo/RNA-free dimer interface.

The domain organization of IMP2 dimer revealed by the SAXS model also enabled us to predict the dimerization interface from the clear domain boundaries. We find an unambiguous head to tail orientation of molecule-1 to molecule-2. The last 12-15 residues from the C-terminus, which forms the terminal helix of the KH4 domain, along with a terminal tail, interact with the first 5-10 residues of the extreme N-terminus as the beginning of RRM1 in the other molecule. IMP2 sequence suggests an abundance of charged amino acids at the termini, enabling ionic interactions to stabilize the interface. From this model of IMP2, it is expected that the dimer interface is created by the primary interaction amongst KH34^1^: RRM1^2^ domains, more specifically α helix interactions between VKQQE and ELHGK residues (**Fig. 1D**). RRM1^1^ and RRM2^1^ make contact in the dimer with interacting residues between a β sheet (YIGNLSPAVT) and a linker (ENVEQVNT) from the respective domains. In the absence of KH12^1^ is central to the stabilization of this dimer, as it forms contacts with RRM1^1^, RRM2^1^, and KH34^1^. KH12^2^ is also observed to be interacting with the RRM2^2^ but not with RRM1^2^. RRM1^1^ contacts KH34^1^ and KH12^1^, while KH34^2^ is seen to be interacting with both RRM1^2^ and RRM2^2^.

Interestingly, the individual monomers within our best-fitting dimeric SAXS model of IMP2 appear vastly different from the AlphaFold predicted model of IMP2 monomers. While SAXS modeling allowed us to approximate the domain interfaces globally, an orthogonal technique was required for residue-level resolution. We performed hydrogen-deuterium exchange mass spectrometry (HDX-MS) to test whether the aforementioned regions that are expected to be interacting within the dimer are susceptible to disruption by a high salt buffer condition (1M NaCl). Multiple regions across constituent domains exhibited significantly increased deuterium uptake in the high salt condition relative to low salt, alluding to a change in protein conformation, with the possibility of the disruption of oligomeric/dimeric IMP2 states into smaller forms (**Fig. 2A**). Overall, HDX-MS provided excellent peptide sequence coverage of our protein, excluding only minor patches. Uncovered spans of amino acids included ∼95-104, ∼160-190, ∼255-259, and ∼333-340 could be missing due to tightly bound RNA blocking access by proteases used in the HDX-MS workflow. The HDX results indicate that changing ionic strength affects primarily the KH1 and KH4 domains, which is in alignment with the protein interface predicted from our dimer model (**Figs. 1D, 2A**). Residues 5-10 from the N-terminus of RRM1 exhibited an 8% increase in deuterium uptake in the 1 M NaCl sample relative to 50mM. RRM2 also exhibited a 6% increase in ∼5 residues at its C-terminal end. KH1 experienced an increase of 5% deuterium uptake in approximately half of the C-terminal-end, which spanned into the KH1-2 linker region. A region of ∼30 amino acids in the middle of the KH2-3 linker experienced 4-5% increase in deuterium uptake in high salt condition, thus more surface exposed to the solvent, alluding to the presence of at least another interactive or closed state involved with that linker region. KH3 has a minor ∼5 residue region that showed 5% increased deuterium uptake at its N-terminal end in high salt condition. The end of the KH3-4 linker was enhanced in deuterium uptake by 7-8% and continued the change almost throughout the entire KH4 domain.

**Figure 2.**
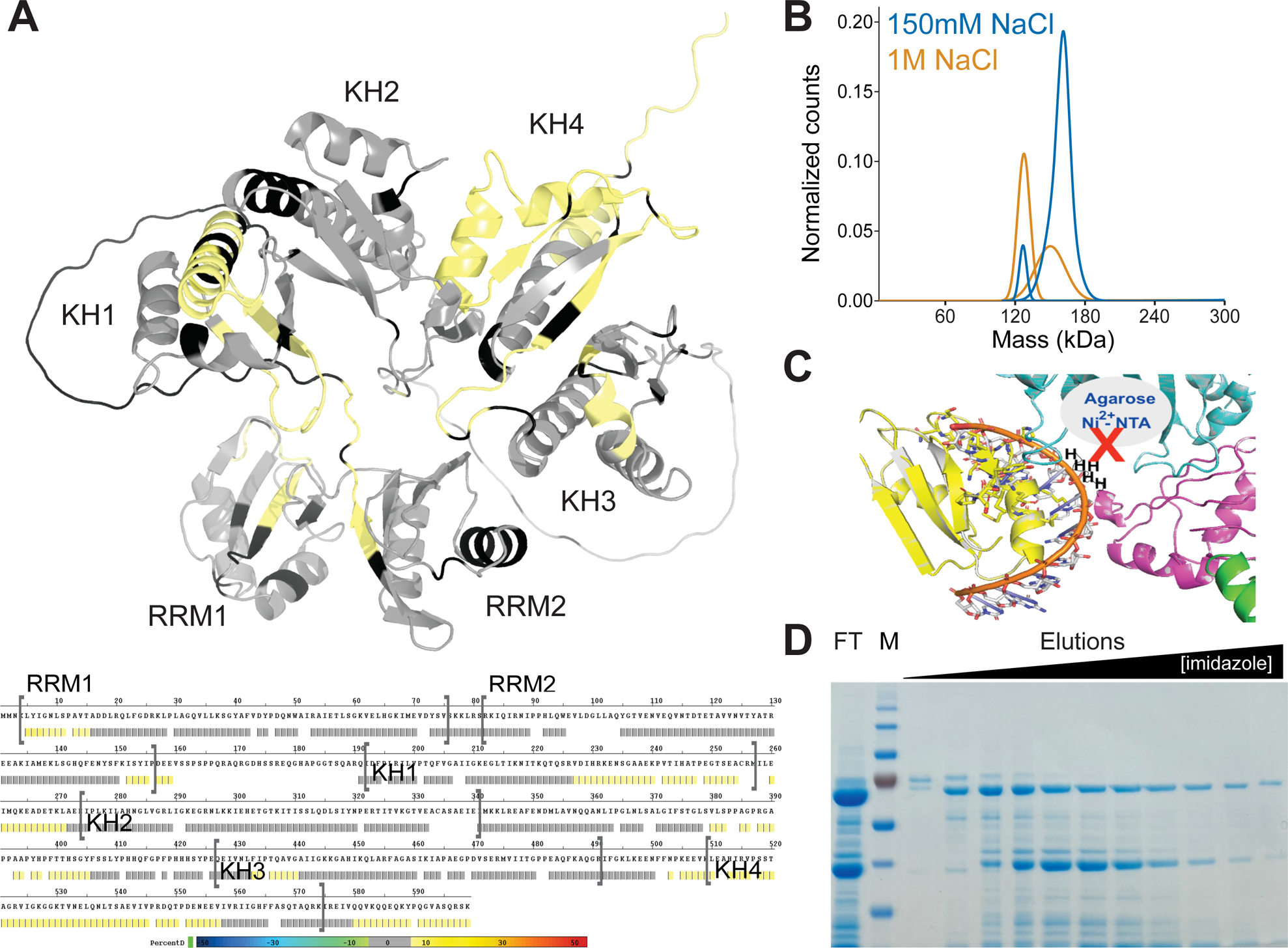
Salt concentration influences IMP2’s architecture. **A)** HDX-MS average deuterium incorporation change of 50mM NaCl vs 1M NaCl mapped onto protein sequence and AlphaFold structure. **B)** Mass photometry of apo/RNA-free IMP2 at 150mM NaCl vs 1M NaCl. **C)** Cartoon illustration of IMP2 highlighting a buried N-terminally placed His-tag. **D)** SDS PAGE gel of His-trap nickel column IMP2 protein purification demonstrating a large amount of protein remaining in flow thought, suggesting the protein was unable to bind adequately in low salt conditions (150mM NaCl). FT-flow through. M – marker.

Mass photometry (MP) confirmed that increased salt concentration in the buffer results in shift from predominantly dimerized form of 160kDa (82% of the protein; monomer size expected as 66kDa; 160kDa species corresponds to a dimer with residual RNA bound during purification; **Fig. 2B and Table S1**) to an even distribution between ^∼^120kDa and ^∼^160kDa species (48% and 42%, respectively). Circular dichroism (CD) spectroscopy also provided complementary evidence that high-salt buffer reduces the stability of IMP2, as the temperature-dependent unfolding profile if IMP2 was reduced from 50 °C in low salt (150mM NaCl) to 41 °C in high salt (1M NaCl) condition (**Fig. S2**). This also suggests the disruption of equilibrium where oligomeric species shift into smaller isoforms. HDX-MS results also reveal a congruent change with easier uptake of deuterium in both previously discussed terminal patches of the protein (**Fig. 2A**). This ‘head-to-tail’ interaction provided an explanation for the better binding of our his-tagged protein to the Ni-affinity columns in presence of high-salt, which would be inaccessible in low ionic-strength buffers (**Fig. 2C and D, Fig. S1A**). Moreover, zeta potential assessment also demonstrated the change in total protein surface charge from approximately −3 at low salt to +5 in high salt exposing several charged residues at the surface (**Table S2**). This could have direct implications in RNA binding as the net positive charge would indicate more residues that can favorably bind negatively charged RNA in such a conformation.

### Structural rearrangements in IMP2 dimer upon RNA binding

To determine how RNA binding influences the structure and oligomeric properties of IMP2, we studied two 67-nucleotide RNA sequences, Cox7b and VSV, that we used previously as an IMP2 binder and non-binding control, respectively, in proximity ligation *in vitro* experiments^3^. Three species of shorter RNA (∼25mer) were recently utilized in a IMP2 small molecule discovery efforts(17); based on this report, we conducted experiments with shorter RNA fragments including two RNA species shown to bind to IMP2, a methylated species (m6A RNA) and an unmethylated IMP2 target (RNA B), as well as a non-binding control (RNA C). mRNA methylation has been recently proposed as an additional level of transcriptional control of gene expression and RNA methylation-dependent binding to IMP2 was shown to affect RNA stability and storage (20, 21).

Since our HDX results indicate the high ionic-strength buffers can induce IMP2 protein conformational changes, we posited that addition of RNA would have an impact on its dynamics or the domain architecture. In general, the IGF2BP protein family is known to bind to RNA and various studies suggested that multiple domains are involved in this interaction (5, 22). To identify the protein surfaces involved in IMP2-RNA interactions, we performed HDX-MS on IMP2 in the absence of presence of Cox7b RNA, a known biologically relevant binding target of IMP2, and VSV RNA, used as a control in *in vitro* experiments (8). Addition of Cox7b RNA (67nt) resulted in occlusion of a multitude of residues throughout the entire length of the protein (**Fig. 3A**). Specifically, a majority of RRM1, RRM1-2 linker, RRM2, KH1-2 linker, KH2, KH3, KH3-4, and KH4 showed a significant decrease in deuterium uptake upon addition of RNA, supporting cooperation between multiple domains in RNA binding. Interestingly, the entire KH1 and a large portion of KH4 showed no peptide coverage in the mass spectrometry experiments, possibly explained by tight ionic interactions between domains of one or multiple IMP2 proteins or even tight interactions between RNA and the protein. This is particularly interesting as these areas exhibited peptide coverage in the low vs high salt HDX experiment, suggesting that this is likely a biologically relevant result and not a technical issue with the digestion or mass spectrometry experiment.

**Figure 3.**
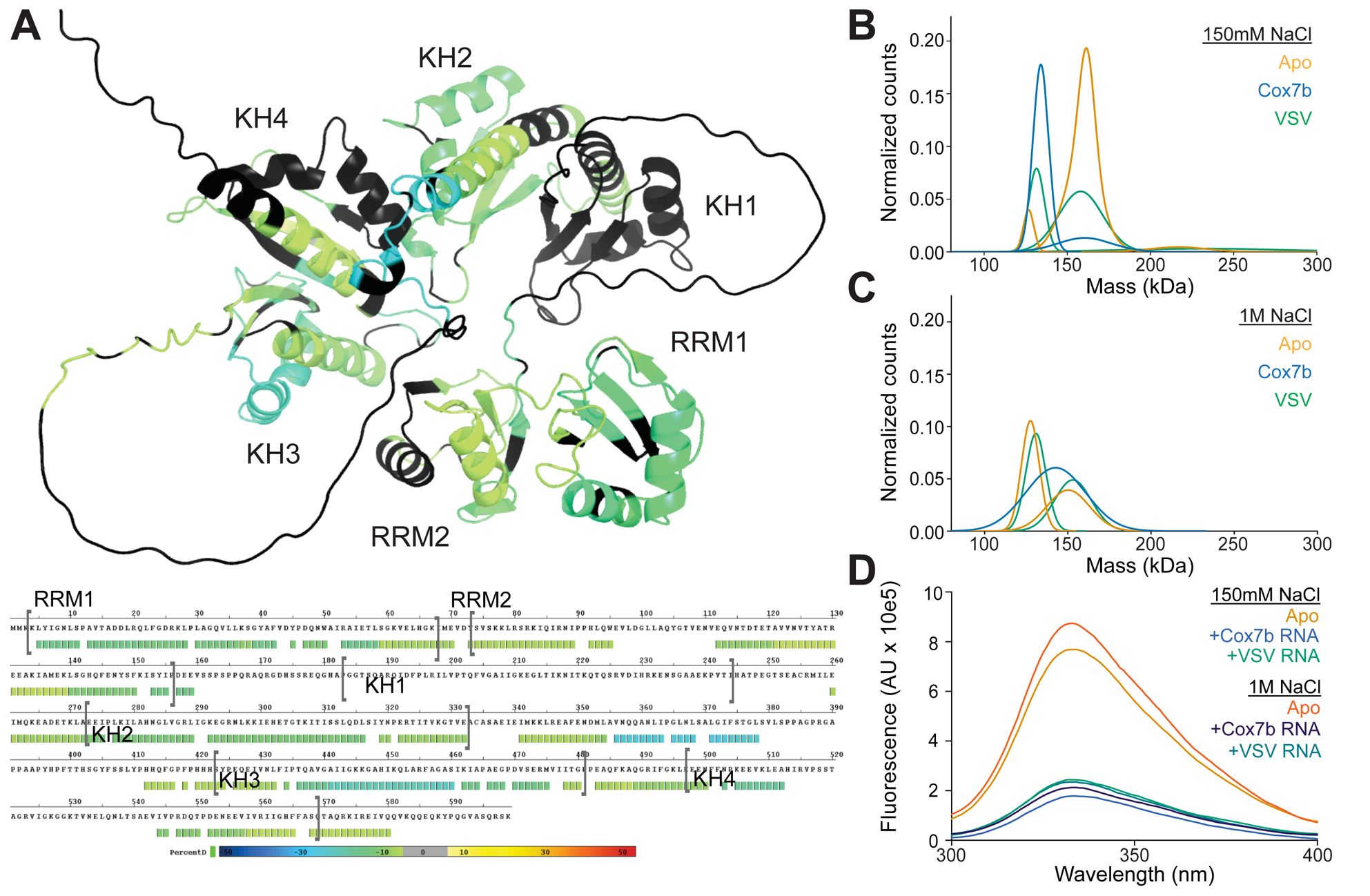
RNA binding alters protein architecture and multimeric equilibrium. **A)** HDX-MS average deuterium incorporation change with vs without Cox7b RNA mapped onto protein sequence and AlphaFold structure. **B)** Mass photometry of apo/ RNA-free, IMP2 + Cox7b RNA, and IMP2 + VSV RNA in 150mM NaCl. **C)** Mass photometry of apo/RNA-free, IMP2 + Cox7b RNA, and IMP2 + VSV RNA in 150mM NaCl. **D)** Fluorescence quenching assay, excitation at 280nm.

To further investigate how addition of RNA could impact IMP2 oligomerization, we performed MP, with the Cox7b and VSV RNAs. The presence of RNA affects the conformation of IMP2 (**Fig. 3B, Table S1**). A multimeric equilibrium appears to be present, which is sensitive to salt concentration (**Fig. 3C, Table S1**). We also observed differences in the UV absorbance for higher order oligomers during SEC-MALS suggesting potential perturbation of dimeric accessibility of surface tyrosines and tryptophans that differs from RNA bound or monomeric forms (**Fig. 3D** and **Fig. S1**, **Table S1**). To test this, we used a fluorescence quenching assay to measure changes of tyrosine and tryptophan fluorescence emissions in IMP2 samples incubated with or without RNA and either a high or low salt solution. There is a drastic change in tyrosine signal reduction for all RNA conditions vs the apo/RNA-free condition (**Fig. 3D**), while tryptophan emissions remain largely unchanged (**Fig. S3**). This supports our hypothesis of conformational change induced by RNA addition and the involvement of tyrosines at the involved interfaces.

To determine if RNA binding influences IMP2 dimer structure, we calculated 3D molecular envelope or shape of Cox7b RNA-bound IMP2 using in-line SEC-SAXS (**Fig. 4A-D**). The RNA-bound IMP2 SAXS data reported a molecular size consistent with an IMP2 dimer; however, the molecular envelope of the RNA-bound IMP2 dimer was more compact than the apo/RNA-free IMP2 dimer (**Fig. 4A** and **B**). Theoretical scattering of the RNA-bound IMP2 SAXS-determined model agreed well with the experimental scattering data (*χ*2 value of 4.99) (**Fig. 4A**). Interestingly, the dimeric form observed in the presence of RNA showed significant differences in the 3D organization of the constituent domains (**Fig. 4B** and **E**). The primary difference is in the dimerization interface, which is formed by the KH12 domains from each of the molecules, yielding a pseudo-symmetric configuration. In the presence of RNA, the IMP2 still predominantly exists in the dimeric form, but the IMP2 conformation is different. The protein-only fit (without RNA) model resulted in a *χ*^2^ value of <10, which indicates a highly concordat fit between the theoretical scattering profile and experimental data. We originally hypothesized that the Cox7b RNA existed in a structured state and bound IMP2 in such a manner. However, upon the addition of every prediction-based folded RNA (RNAfold web server) individually into the SAXS model, the *χ*^2^ value worsened greatly indicating a poor fit and an unlikely model (**Fig. 4A**). This suggested two possibilities: that the RNA does not bind to protein in our experimental conditions or the RNA binds in an unstructured manner. IMP2 interaction with RNA helicases also supports the possibility that the target RNA bound to the protein might be in an unfolded state (8, 23). Furthermore, there is little to no space in the calculated SAXS envelope to fit the folded RNA in once the protein portion is fitted. We therefore added the RNA in an unstructured fashion, which resulted in *χ*^2^ value 4.99, suggesting that IMP2 dimer may interact with unstructured RNA.

**Figure 4.**
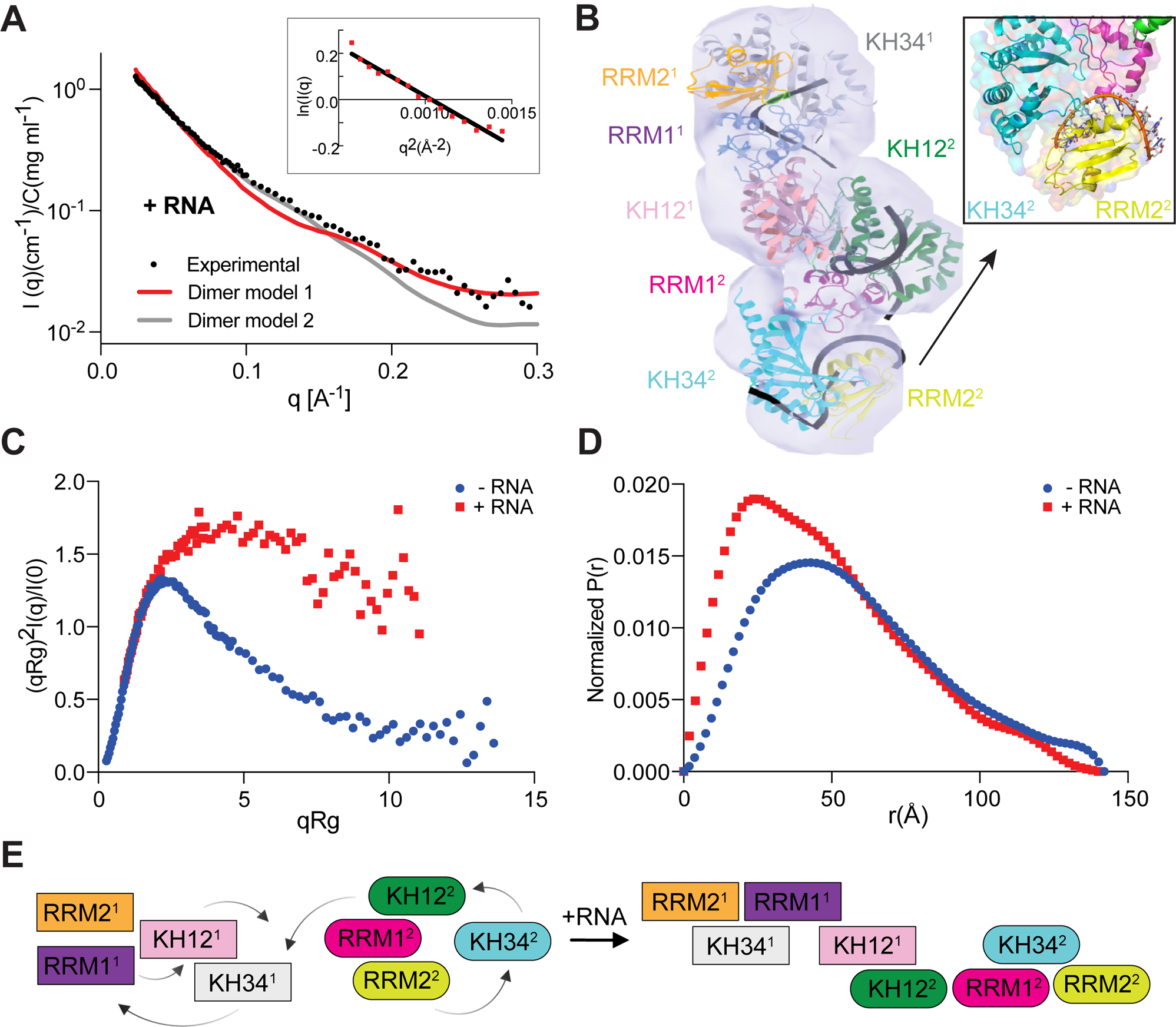
Apo/RNA-free and RNA-bound IMP2 dimers are unique. **A)** The log I(q) versus q plot represents the primary SAXS data, with Guinier plots shown as insets. The maximum dimensions RNA-bound IMP2 and apo/RNA-free IMP2 are 142 and 140Å respectively, and the minimum q measured (0.006 Å^−1^) is well below the minimum recommended for accurate assessment. In both experiments, the Guinier plots (inlay) are linear to the first measured q values and a plot of log I(q) versus log q shows that the slope is effectively zero at low q as expected for monodisperse particles of similar size. These measures together provide confidence that the data are free of significant amounts of contaminating species or inter-particle correlations contributing a structure-factor term to the scattering. **B)** SAXS envelope with RNA-bound dimer model within. Inlay of zoomed in RNA-protein interaction. **C)** Dimensionless Kratky plots demonstrate that the SAXS data are from predominantly folded particles. The IMP2 +/− RNA plots display the expected bell-shaped curves, with a maximum of about 1.3 at around qR_g_ = 2.07 for the apo/RNA-free condition and maximum of 1.7 at around qR_g_ = 4.78 for the apo/RNA-free condition. The RNA-bound complex is significantly more elongated rather than being globular, as observed from the shallow tapering of the Kratky plot. The peak for IMP2 RNA-bound is slightly shifted to the right as expected for its slightly elongated shape, and the small rise of apo/RNA-free IMP2 evident at qR g > 11 suggests some flexibility. **D)** The P(r) versus r profiles for each of the proteins are well behaved, showing the smooth, concave approach to zero at r = 0 and D_max_ expected for a mostly folded, monodisperse protein. IMP2 apo/RNA-free has a D_max_ of 142 while IMP2 RNA-bound has a D_max_ of 140. For both conditions, the R_g_ and I(0)-based M values are in excellent agreement between independent Guinier and P(r) analyses. **E**) Schematic representation of the domain changes from apo/RNA-free dimer to RNA-bound dimer.

Our 3D reconstruction of the dimeric IMP2-Cox7b RNA-bound complex shows a more compact conformation and a global alteration of the protein conformation compared to that of IMP2 dimer in the apo/RNA-free form. The structural models indicate that RNA-bound dimeric IMP2 may be more stable and thermodynamically favorable with a reduction of the number of accessible surface atoms (apo/RNA-free 4363 vs RNA-bound 3976) and increase in the number of buried atoms (apo/RNA-free 2805, RNA-bound 3192). The radius of gyration (R_g_) of IMP2 dimer decreases upon binding of RNA from 46.1 ± 1.2 to 36.8 ± 1.6 Å, suggesting a more compact complex. At the same time, the slight decrease in the D_max_ value (142 ± 1 vs 140 ± 1 Å) is observed, meaning that the compression is not uniform but instead unidirectional (**Fig. 4D**). Dimensionless Kratky plots (**Fig. 4C**) indicate that RNA-bound IMP2 is significantly more elongated as opposed to globular, as observed from the shallow tapering of the curve, typically found in complexes with both folded and unfolded elements. In our case the RNA is likely unfolded, leading to a composite of folded and unfolded features. When comparing protein in the presence of RNA against its dimer in the absence of RNA, the apo IMP2 dimer exhibits a head-to-tail configuration, with RRM1 and RRM2 at the terminal end of one IMP2 and KH34 at the dimer interface and KH12 in between (**Fig. 4E**). The RRM1^2^ appears to interact with KH34 of IMP^1^. KH12 and RRM2 are located between RRM1 and the terminal KH34. Upon binding of RNA, RRM1 from IMP2^1^ moves from the terminal end of molecule 1 to the interior closer towards the dimer interface near molecule 2 (by ∼ 23Å), while position of RRM2 remains relatively unchanged. KH12 switches position with KH34, such that in presence of RNA it is located between RRM1/RRM2 and the newly formed KH12 interface. IMP2^2^ also exhibits the relative reorganization of its KH12 domain, positioning itself more interiorly, forming a new KH12-KH12 interface. KH34 moved internally from the terminal end of the second IMP2, with RRM1 and RRM2 migrating exteriorly.

While there is a global conformational change between the apo/RNA-free and RN-bound dimers, some domain interactions are consistent in the structural models. RRM1^1^ and RRM2^1^ make contact in both dimers with the modification of contact residues from amino acids 6-15 : 109-116 (β strand and linker) to 15-33 + 68-73 : 82-89 + 116-124 + 152-157 (Alpha and linker interactions). Similarly, RRM2^2^ and RRM1^2^ remain in proximity and retain α helix and linker contact (amino acids 139-146 + 153-157 : 29-34 + 51-62 vs 132-142 : 3-9). This differs greatly from RNA-bound dimer KH12^1^ which has little to no contact with RRM2^1^ and KH34^1^ but remains in proximity to RRM1^1^. Contact between KH12^2^ and RRM2^2^ exists in the apo/RNA-free dimer, but not the RNA-bound dimer. While RRM1^2^ and KH12^2^ do not interact in apo/RNA-free dimer, they interact in RNA-bound dimer. In contrast to the extremes of total reconfiguration or no change, certain domains (for example RRM2^2^-KH34^2^, an α helix vs β sheet involvement) remain in contact, yet in an alternative configuration.

The apo/RNA-free IMP2 dimer interface is created by the primary interaction amongst KH34^1^: RRM1^2^ domains, more specifically α helix interactions. Despite the critical nature of these domain interactions in apo/RNA-free dimer, the RNA-bound dimer exhibits no interaction amongst these domains. The domain interface of Dimer 2 is instead driven by KH12^1^ and KH12^2^ domains (β sheet interactions between amino acids 193-203 + 224-243 + 313-320 and 223-247). It is clear that a major structural change occurs in the presence of RNA with two distinct interfaces with different underlying interactions (α helix vs β sheet). Furthermore, the interactions amongst KH34^1^: RRM1^2^ only exist in apo/RNA-free dimer while the interactions amongst KH12^1^ and KH12^2^ are unique to RNA-bound dimer.

Our HDX-MS data supports this observation as a high ionic-strength environment exposed several residues from the KH12 domain upon exposure to a buffer containing high concentration of NaCl (1M) suggesting the disruption of the KH12-KH12 interface in the RNA bound dimer, exposing them to deuterium uptake. Additionally, KH34 and RRM1 also show areas of enhanced deuterium uptake in presence of high ionic strength, supporting the opening of the unbound dimer interface. Despite changes in deuterium uptake at the newly exposed surfaces of IMP2 domains, some regions experience no change (for example, residues ∼20-150, ∼190-225, ∼270-375, ∼440-500), ensuring the protein to be folded and not denatured by the high ionic-strength environment. Most of the core regions of the domains of IMP2 remain unaltered, suggesting that the changes associated with the protein are predominantly coming from the conformational rearrangement of the domain interfaces. RNA binding to IMP2 is also observed to involve almost the entire full-length protein (evident from the changes in the buried residues of IMP2 in HDX), except for a significant portion of KH12 domains, which is in line with SAXS modeling with KH12-KH12 interface being formed in presence of RNA.

### Insights into the mode of RNA binding

Upon placement of the Cox7b RNA in five unfolded fragments into the SAXS envelope, we noted a marked improvement of the *χ*^2^ values. Further inspection reveals that the dimerization interface is critical to ensure the RNA interacting groves are exposed towards the substrate nucleotides. Major interaction surfaces are provided by the last α helix and portions of the linker region of the RRM2^2^ domain to interact with nucleotides of Cox7b RNA (**Fig. 4B and E**). While it is impossible to determine the exact residues involved in RNA-protein interaction from SAXS, we notice from our tyrosine quenching studies (**Fig. 3D**) that the FENYSF stretch from the RRM2^2^ domain and LYIG from the RRM1^2^ domain could be involved in the π-π stacking interactions stabilizing the RNA. Another RNA fragment is observed to be positioned at the central grove of the KH12 domain and likely to be interacting with the SIYNPER loop within the domain (**Fig. 4B**). It is important to note that the backbone phosphates from the RNA are also likely to have favorable water mediated hydrogen-bond interactions with the terminal residues of the helices, as the resulting dipole moments from them will allow for increase in the electronegativity of the participating atoms, which are well aligned for such interactions.

The combined evidence from tyrosine quenching assays, HDX and the envelope from SAXS strongly suggest a proteinaceous ‘cocoon’ (**Fig. 4B**), where most of the RNA nucleotides are protected from solvent interactions. Upon examination of the surface charge of the structural model, we also notice a significantly large, connected patch of negatively charged residues throughout the second molecule of the dimer and another mildly positive surface on the other side of the dimer (**Fig. S4**). These two surfaces create a unique sheet like architecture with one face being particularly negatively charged and the other being positive along the long edge of the dimer.

While there is a change in RNA binding based on the length and nature, the RNA always exhibits interaction with IMP2 throughout our experiments. This supports the notion that the binding does not discriminate based on specific base-pairing nor RNA structure. Fluorescence polarization (FP) assays performed at physiologic salt concentration (150mM NaCl) with shorter RNA oligonucleotides previously tested for IMP2 binding (17, 20, 21) show the highest binding affinity to RNA B (IMP2 binder), next to RNA m6A (m6A modified IMP2 binder), and least affinity to control RNA C (**Fig. 5A**). Strength of charge-dependent protein-protein and protein-RNA interactions typically decreases in high salt buffer conditions. The RNA affinities for IMP2 in our FP experiments differ when performed with high salt (1M NaCl), with RNA m6A becoming the highest affinity, RNA B second, and then the control RNA (**Fig. 5B**). As we suspect the dimeric IMP2 to be breaking in high salt condition, this could allow for newly exposed protein regions to interact with RNA, which is in line with our HDX and SAXS data. However, in the presence of low-salt buffers there are multiple species, while high salt appears to bias for one species. Despite the change in affinity, the RNA is still capable of binding to the protein in the presence of high salt, showcasing the strength at which RNA binds to the protein. This is supported by our SEC-MALS results which show the disruption of multimeric IMP2 states into the smaller isoforms while still allowing RNA to remain bound (**Table S1**).

**Figure 5.**
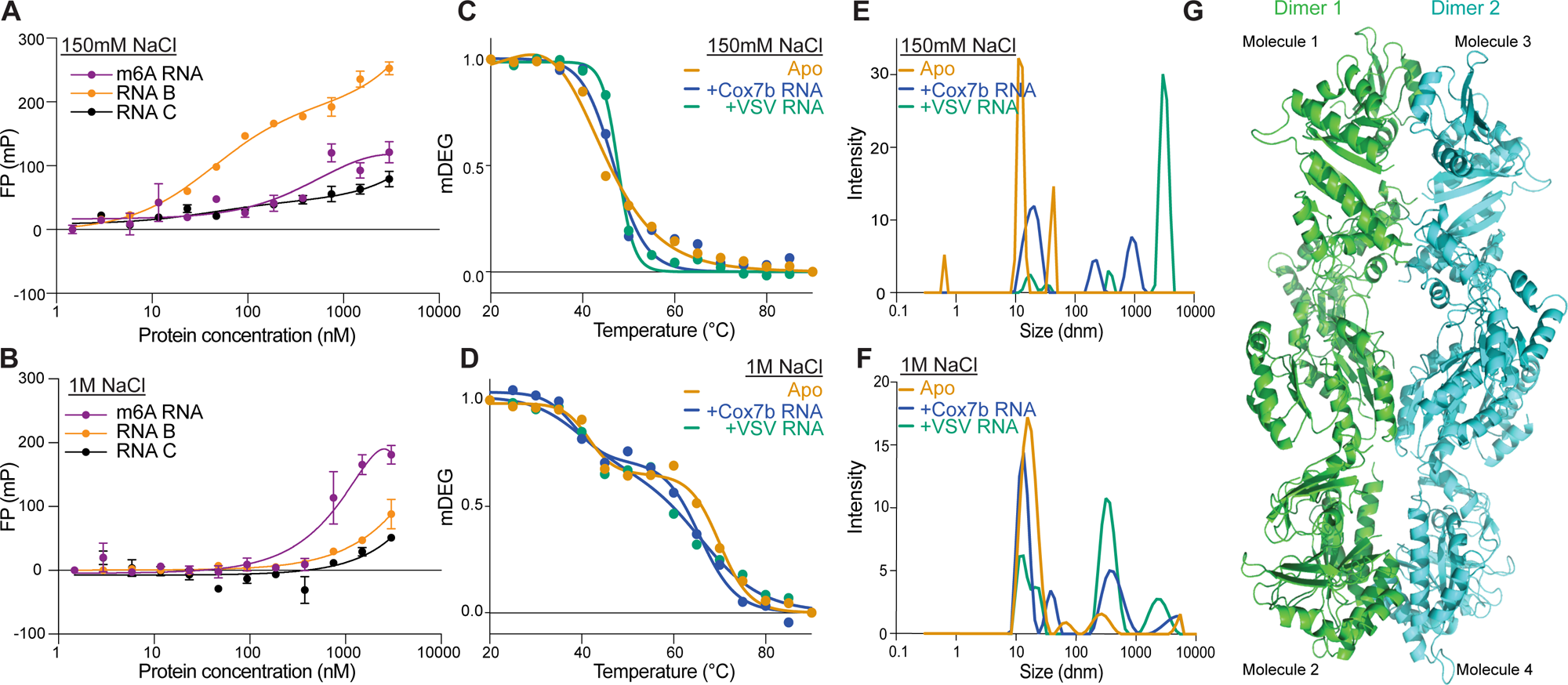
RNA species impacts IMP2 properties. **A)** Fluorescence polarization of IMP2 and fluorescently labeled RNAs in 150mM NaCl buffer. **B)** Fluorescence polarization of IMP2 and fluorescently labeled RNAs in 1M NaCl buffer. **C)** CD thermal denaturation of apo/RNA-free (T_m1_ = 50°C), IMP2 + Cox7b RNA (T_m1_ = 47°C), and IMP2 + VSV RNA (T_m1_ = 44°C) in 150mM NaCl buffer**. D)** CD thermal denaturation of apo/RNA-free (T_m1_ = 41°C, T_m2_ = 70°C), IMP2 + Cox7b RNA (T_m1_ = 39°C, T_m2_ = 66°C), and IMP2 + VSV RNA (T_m1_ = 40°C, T_m2_ = 66°C) in 1M NaCl buffer. MALS of apo/RNA-free, IMP2 + Cox7b RNA, and IMP2 + VSV in 150mM NaCl buffer. **F)** MALS of apo/RNA-free, IMP2 + Cox7b RNA, and IMP2 + VSV in 1M NaCl buffer. **G)** Model of “dimer stacking” with two RNA-bound dimer models.

As mentioned above, VSV RNA has been previously demonstrated as a non-binding RNA in a cellular context (8). However, in our electric mobility shift assay (EMSA) VSV RNA appeared to bind IMP2, similar to Cox7b RNA (**Fig. S5**). There are several potential explanations, including but not limited to oligomeric states of IMP2, protein-protein interactions and/or posttranslational modifications that are likely modulating specificity of target binding. Interestingly, we see the alteration of RNA binding affinity in different environments in our FP assays. Altogether, these results underscore the importance of the environmental and modular arrangements in which IMP2 can coordinate and enhance binding to different RNA targets.

CD thermal denaturation curves of IMP2 +/− RNA samples exhibited unique profiles depending on the salt concentration. Samples exposed to low salt (150mM) exhibited a classic 2-state profile, starting from a native/folded state and proceeding to an unfolded/denatured state as temperature is gradually increased (**Fig. 5C and D**). Interestingly, the exposure of these samples to high salt (1M) greatly altered the thermal denaturation pattern and displayed an “intermediate” state as evidence by its biphasic curve/3-state profile (**Fig. 5D**). While apo/RNA-free IMP2 in 150mM NaCl has Tm_1_= 50°C, the addition of RNA in the same buffer system appears to decrease the melting temperatures (VSV: Tm_1_=44°C; Cox7b: Tm_1_=47°C). In high salt buffer (1M NaCl), the Tm_1_ for apo/RNA-free IMP2 (41°C), Cox7b + IMP2 (39°C), and VSV + IMP2 (40°C) are similar, while the Tm_2_ of RNA-bound IMP2 (66°C) conditions are approximately 4°C lower than that of apo/RNA-free IMP2 sample (70°C). These results are counterintuitive at first glance because we anticipated that binding of RNA would provide enhanced protein stability and encourage the formation of oligomers. While the IMP2 + RNA conditions show a decrease in Tm_1_, between low and high salt conditions, the Tm_2_ increases. We suspect that high salt could break IMP2 oligomers into less stable smaller isoforms. While protein thermal unfolding is irreversible, a 3-state unfolding appears to be present with an intermediate existing between folded and unfolded states. This intermediate could indicate a dissociation event, either between the oligomers of the protein or between the protein and its RNA partner. MALS results also confirm that binding of two different RNA probes results in shifts towards higher order oligomers, which are sensitive to the presence of high salt (1 M NaCl; **Fig. 5E and F**).

#### Higher Order Oligomers of IMP2

Our results indicate that IMP2 interacts with RNA in a dimeric form that is distinct from its apo/RNA-free dimer. Our models suggest that the head-to-tail positioning of the Apo/RNA-free IMP2 dimer rearranges to form a more symmetric dimer in the RNA bound state. Furthermore, RRM2 domains move to be positioned at the terminal ends of the elongated dimer structure, with KH34 and RRM1 domains between the terminal RRM2 and internal KH12 domains. The interface of interactions was stabilized by KH12-KH12 domains in the RNA bound form vs RRM1 and KH34 domains in the unbound state. Interestingly, when two IMP2 dimers are docked to each other to generate a tetramer, complementary charged surfaces form a stacked-dimer orientations (**Fig. 5G**). Out of the 200 possible docked poses, the analysis was continued with best scoring model. This docked model appears to be of the most thermodynamically favorable among other possibilities which we cannot rule out based on *in silico* assessments only. The prevalence of such charge-based interaction could be the reason why in a high ionic-strength environment, the oligomers dissociate into a stable dimer, which is not susceptible to rupture under such conditions.

Examination of the docked tetrameric structure allowed us to find that the two dimers, both in the RNA bound dimer configuration, come together to form a dimer stacking tetramer. This tetramer or ‘stacked-dimer’ of dimers is maintaining a parallel orientation, such that the molecule 1 from dimer 1 is located just on top of the molecule 1’ or dimer 2. The ‘stacked-dimers’ only partially use the extensive negatively charged surface, allowing for other partners to bind, maybe even other molecules of IMP2 (**Fig. 5G**). Core of this structure is the most central contact point containing the convergence of all KH12 domains from both dimers (2x KH12^1^ and 2x KH12^2^ domains). Other contact points between the pair of dimers include RRM1^2^, RRM2^2^, and KH34^1^ domains. External, non-contacting domains consist of RRM1^1^, RRM2^1^, and KH34^2^. Furthermore, the “sandwiching” effect of stacking dimers could further sequester RNA, in line with the difficulty of removing RNA from IMP2 purified proteins. This stacking type mechanism could also provide further clues into RNA protective effects that RNA binding proteins typically exhibit in the context of cellular stress (stress granule formation, etc.) (24).

While many possible higher order multimeric states of IMP2 appear to exist, as evidenced by our MP, SEC-MALS, and SAXS data, they seem be multiples of two (**Table S1**). This could suggest that IMP2 dimers are the predominant building block, resulting in higher order oligomers. In addition to the higher order multimers, we hypothesize that some of the larger complexes could participate in stress granule or condensate formation as a protective cellular mechanism to sequester RNA and prevent degradation in cellular stress situations, as it has been shown for other RNA-binding proteins (25). Interestingly, we see an alteration in UV absorbance for higher order oligomers during SEC-MALS which could suggest perturbation of the solvent accessibility of tyrosines and tryptophans which differs from the RNA-bound states. This also explains why we faced several challenges trying to decouple the native RNA during our recombinant purification, to the point that it maintained a lot of RNA-complex even in presence of an RNase or in presence of LiCl, which is a known RNA precipitating agent.

## DISCUSSION

With a combination of SAXS, HDX-MS, MP, MALS, and SEC-MALS we demonstrate that IMP2 exists as multiple states in solution, with the predominant form being a dimer. Intriguingly, the dimeric species of IMP2 can vary greatly in structural organization of its domains when comparing the conformation of RNA bound and unbound states. The apo/RNA-free IMP2 dimer exhibits a head-to-tail positioning with KH34 and KH12 at the interface, while the RNA-bound dimer adopts a more symmetric configuration stabilized by a KH12-KH12 interface. Previous studies have demonstrated the capability of IMP1, IMP2, IMP2 splice variant, and IMP3 to form homodimers in the presence of an RNA target (13, 26). These full-length proteins also formed heterodimers with IMP1’s KH1-4 domains (13). Interestingly, only IMP2 and IMP2 splice variant appeared to form higher-order complexes larger than the apparent dimer (13). Our data supports the notion of higher molecular weight IMP2 multimers and these various IMP2 states appear to be sensitive to changes in ionic strength. Since IGF2BP proteins are known to participate in membraneless organelles, such as stress granules (24, 25), the sensitivity of oligomerization to changes in ionic strength could reflect functional response to local changes in environmental conditions *in vivo*, allowing for fast regulation of stability of RNA targets of IMPs.

Our HDX-MS data suggest that all IMP2 domains are involved in binding of our 67nt Cox7b RNA probe. The addition of Cox7b RNA also altered the relative abundance of IMP2 species detected by MP, changed the CD thermal melt profiles, and even resulted in the quenching of tyrosine intrinsic fluorescence. Previous studies also show that IGF-II RNA encourages the formation of homo- and heterodimers amongst IMP paralogs and that H19 RNA interacts with four molecules of IMP1(13, 27), supporting the RNA-driven oligomerization. In order for this RNA species to interact with the entirety of IMP2, it is likely that the RNA secondary structure becomes unfolded. Our SAXS data support this hypothesis as the χ^2^ value is worsened when fitting the RNA as structured molecule, while modeling the unstructured RNA improves the fit. Recent literature proposed unstructured and unaided RNA binding as a possible mode of IMP-RNA interaction (22). Moreover, RNA helicases have been identified as interacting partners of IMP3 and IMP2 (8, 28), adding to the possibility of unfolded RNA binding by IMP proteins.

Intrinsically disordered and/or linker regions are known to be important for multiple protein functions but pose a major challenge to structural studies of full-length proteins (1, 29, 30). Our data suggest that protein-protein and protein-RNA interactions occur among both IMP2 structured domains and unstructured linker regions. Further biological relevance and importance of these linker regions is exemplified by studies demonstrating how condensate formation is impacted by as little as a single amino acid modification positioned in these unstructured stretches (31). Especially in the context of IGF2BPs, these unstructured protein elements appear to be critical for RNA binding specificity, protein domain organization and multimeric state interactions (5, 32).

By utilizing a combination of SAXS and HDX-MS, we uncovered domain level information on full-length IMP2 in RNA bound and unbound dimeric states. To obtain atomic resolution structural data, X-ray crystallography or cryo-electron microscopy (cryo-EM), methods which require a homogenous protein sample, can be leveraged. By understanding the dynamics and influences of multimeric equilibrium, our data indicate that future studies focused on optimizing a homogenous IMP2 sample preparation may improve the odds of using these techniques. Moreover, the comparison of purified protein studies to *in vitro* experiments are necessary to ensure that what we observe is biologically relevant and applicable. Understanding the intricacies of protein-protein and protein-RNA interactions, and their sensitivity to external influences and protein/RNA partners will be critical to fully understand the molecular basis of IMP2 function which are key factors to consider in IMP2 drug discovery efforts in cancer and metabolic disease.

## EXPERIMENTAL PROCEDURES

### Protein purification

Full length IMP2 protein was expressed as a pET-45 6-Histidine tagged 3C protease cleavable protein in Rosetta (DE3) pLysS *Escherichia coli* (Novagen) cells at 18 °C overnight; the expression was induced with 0.5 mM IPTG. Bacteria were lysed, and the protein was isolated on a HisTrap HP nickel−sepharose column (Cytiva) followed by an imidazole buffer elution. Cells were harvested by centrifugation at 4000g and resuspended in binding buffer (20 mM HEPES, pH 7.4, 2 mM MgCl2, 150 mM NaCl, 10% v/v glycerol, 2 mM mercaptoethanol, 15 mM imidazole). Ethylenediamine tetraacetic acid -free protease inhibitor cocktail (Roche) was added fresh to the binding buffer. Cells were lysed using a sonicator, and cell debris was removed by centrifugation at 20,000g at 4 °C for 1 h.The supernatant was applied to a 5 mL HisTrap HP nickel− sepharose column at 1 mL/min on an ÄKTA system. The column was equilibrated with a binding buffer in a 10 × column volume or until the UV signal was stable. The protein was eluted with a linear gradient to 100% of 500 mM imidazole buffer (HEPES, pH 7.4, 2 mM MgCl2, 150 mM NaCl, 10% glycerol v/v, 2 mM mercaptoethanol, 500 mM imidazole) at 1 mL/min. Fractions containing IMP2 were collected and concentrated via centrifugal filtration using Vivaspin columns (30,000 MWCO, Sartorius), resuspended in SP buffer (HEPES, pH 7.4, 2 mM MgCl2, 50 mM NaCl, 10% glycerol v/v, 2 mM mercaptoethanol), and loaded onto a 5mL HiTrap SP column. Protein was eluted with a linear gradient to 100% 1M NaCl buffer (HEPES, pH 7.4, 2 mM MgCl2, 1M naCl, 10% glycerol v/v, 2 mM mercaptoethanol). Pure fractions were collected, concentrated, and ran on an S200 size exclusion chromatography column with SEC buffer (HEPES, pH 7.4, 2 mM MgCl2, 150 mM NaCl, 10% glycerol v/v, 2 mM mercaptoethanol). Protein purity and identity were assessed by SDS-PAGE, Western blot, and CDS. The eluted IMP2 fractions were concentrated via centrifugal filtration using Vivaspin columns (30,000 MWCO, Sartorius). Protein concentrations were measured by both UV spectroscopy (ε = 280 nm) and Bradford assay (according to the manufacturer’s instructions). Protein aliquots were stored at −80 °C.

### RNA

The RNA oligonucleotides utilized in this study were previously described (8, 17). Cox7b_RNA (67nt, Integrated DNA Technologies): 5’- UCAUCCCAGCUGGUGUAAUAAUGAAUUGUUUAAAAAACAGCUCAUAAUUGAUGCCAAAUUAAAGCAC-3’. VSV_RNA (67nt, Integrated DNA Technologies): 5’- ACAUAAAAAGCUUUUUAACCAAGCAAGAAUGAAGUAUCGUAUCUAAUUAAUUCCGAUGAU CAAUAUG-3’. RNA_m6A (20nt, Dharmacon/Horizon Discovery): 5’-Fluorescein-CGUCUCGGm6ACUCGGm6ACUGCU-3’. RNA_B (25nt, Dharmacon/Horizon Discovery): 5’- CCCCCCUUUCACGUUCACUCUGUCU-3’-Fluorescein 3’. RNA_C (25nt, Dharmacon/Horizon Discovery): 5’-Fluorescein-GAAAAAAAGAUUUAUUUAUUUAAGA-3’.

### Buffers

Unless otherwise specified, “low” and “high” salt buffers refer to the following: low salt buffer (20mM HEPES, pH 7.4, 2 mM MgCl2, 150mM NaCl, 10% glycerol v/v, 2 mM mercaptoethanol) high salt buffer (20mM HEPES, pH 7.4, 2 mM MgCl2, 1M NaCl, 10% glycerol v/v, 2 mM mercaptoethanol).

### Fluorescence polarization (FP)

FP assay with 20nt probes was performed as previously described (17). Lyophilized RNA oligomers were dissolved in the low salt buffer. 100 μM stock solutions of RNAs were created which were further diluted into 100 nM aliquots and stored at −80 °C. A constant concentration (1nM final) of each FLC-labeled RNA was titrated in combination with serial dilutions of IMP2 protein (0.15 nM to 3 μM final). IMP2 and FLC labeled RNA were incubated for 1 hour at room temperature in 384-well black microplates before the start of plate reading. Fluorescence polarization and fluorescence intensity were measured using a BioTek Synergy Neo 2 microplate reader with an excitation at 485−495 nm and an emission at 520−530 nm. Focal height and gain adjustments were done before starting each measurement. Each sample was tested in triplicate, with FP values reported in millipolarization units (mP). GraphPad Prism (version 10) was used to plot and fit the data to a one-site binding “total” equation.

### Hydrogen-deuterium exchange coupled to mass spectrometry (HDX-MS)

Differential HDX-MS experiments were conducted as previously described with a few modifications (31). Peptides were identified using tandem MS (MS/MS) with an Orbitrap mass spectrometer (Q Exactive, ThermoFisher). Spectra were acquired in data-dependent mode with the top five most abundant ions selected for MS2 analysis per scan event. The MS/MS data files were submitted to Proteome Discoverer 2.5 (Thermo) for peptide identification using a workflow incorporating Sequest and Percolator. Peptides included in the HDX analysis peptide set were within 10 ppm tolerance to the theoretical molecular weight and classified with high confidence score in Proteome Discoverer. The protein database included the recombinant IMP2 sequence and a list of common contaminants (33). HDX-MS analysis: RNA was prepared from a 100 µM stock in water and diluted into IMP2 in buffer (50 mM HEPES, 10 mM MgCl2, 50 mM NaCl, 1 mM BME, 5% glycerol, pH 7.5) to give a final concentration of 11 µM RNA and 10 µM IMP2. For apo sample, water was used in place of RNA solution. For high salt treatment of IMP2, the sample was prepared analogously to the apo solution above, but with high salt buffer (50 mM HEPES, 10 mM MgCl2, 1 M NaCl, 1 mM BME, 5% glycerol, pH 7.5). After 30 min. incubation (rt), 5 ml of sample then was diluted into 20 ml D_2_O buffer (50 mM HEPES, 10 mM MgCl2, 50 mM NaCl, 1 mM BME, 5% glycerol, pH 7.9), or for high-salt experiment (50 mM HEPES, 10 mM MgCl2, 1 M NaCl, 1 mM BME, 5% glycerol, pH 7.9), and incubated for various time points (0, 10, 60, 900 and 3600s) at 4°C. The deuterium exchange was then slowed by mixing with 25 µl of cold (4°C) 0.1 M sodium phosphate monobasic with 50 mM TCEP. Upon injection, quenched samples were passed through a co-immobilized pepsin/fungal XIII protease column (1mm × 2cm) at 50 µl min^−1^ and the digested peptides were captured on a 1mm × 1cm C_8_ trap column (Agilent) and desalted. Peptides were separated across a 1mm × 5cm C_18_ column (1.9 ml Hypersil Gold, ThermoFisher) with a linear gradient of 4% - 40% CH_3_CN and 0.3% formic acid, over 5 min. Sample handling, protein digestion and peptide separation were conducted at 4°C. Mass spectrometric data were acquired using an Orbitrap mass spectrometer (Q Exactive, ThermoFisher). HDX analyses were performed in triplicate from single preparations. The intensity weighted mean m/z centroid value of each peptide envelope was calculated and subsequently converted into a percentage of deuterium incorporation. This is accomplished by determining the observed averages of the undeuterated and fully deuterated spectra and using the conventional formula described elsewhere (34). Statistical significance for the differential HDX data is determined by an unpaired t-test for each time point, a procedure that is integrated into the HDX Workbench software (35). Corrections for back-exchange were made on the basis of an estimated 70% deuterium recovery, and accounting for the known 80% deuterium content of the deuterium exchange buffer. Data Rendering: The HDX data from all overlapping peptides were consolidated to individual amino acid values using a residue averaging approach. Briefly, for each residue, the deuterium incorporation values and peptide lengths from all overlapping peptides were assembled. A weighting function was applied in which shorter peptides were weighted more heavily and longer peptides were weighted less. Each of the weighted deuterium incorporation values were then averaged to produce a single value for each amino acid. The initial two residues of each peptide, as well as prolines, were omitted from the calculations. This approach is similar to that previously described (36).

### Zeta potential and MALS

The particle size and zeta potential (electrophoretic light scattering) were measured at 25 °C after incubating for 120s using the Zetasizer Ultra Red Label from Malvern Instruments, under various buffer conditions. Disposable folded capillary cells from Malvern were utilized for zetasizer measurements, whereas a low volume quartz cuvette was used for MALS. Zeta potential was calculated using the Smoluchowski model. The data were analyzed utilizing Zetasizer Software to compute the parameters of volume versus hydrodynamic radius.

### Fluorescence spectroscopy

Fluorescence spectra were recorded with a Horiba Fluoromax 3 (Jobin Yvon, Kyoto, Japan) spectrofluorometer. Tyrosines show a maximum of excitation at 280 nm, while typtophans show a maximum of excitation at 295 nm (37). The emission was collected between 300 and 400 nm to probe the resulting changes to IMP2 tertiary structure at different equilibration conditions. The emission spectra was collected after excitation at 280nm and 295 nm, using either IMP2 and RNA alone or a 1:1 ratio of IMP2 and RNA incubated in low or high salt buffer after equilibrating overnight at respective conditions. GraphPad Prism (version 10) was used to plot the data.

### Thermal melt of IMP2 monitored using circular dichroism

IMP2 was buffer exchanged into either low (150mM) or high (1M) NaCl buffer conditions, as previously described. Thermal melts of IMP2 under low and high salt conditions after incubation with (1:1) or without RNA were obtained by monitoring the CD spectra from 200nm to 280nm while increasing the temperature by 5°C from 20 to 95 °C. The signal at 220nm, which shows a significant variation in signal as a function of temperature, was used for further analysis. The signal at a wavelength of 220nm was normalized and plotted for each experimental condition. The Tm was determined by fitting two-state or three-state thermal unfolding equations as previously described by Norma et al. to the data using GraphPad Prism (version 10) (38).

### Size exclusion chromatography coupled to multi-angle light scattering (SEC-MALS)

The purified IMP2 protein +/− RNA was analyzed by SEC-MALS using an HPLC system (Agilent Technologies 1260 Infinity) connected to a MALS system (Wyatt DAWN HELEOS II Ambient with Optilab TrEX HC differential refractive index detector). The SEC–MALS system was calibrated with bovine serum albumin before protein and protein + RNA runs. The gel filtration purified protein was loaded onto pre-equilibrated SEC analytical column (WTC-050S5, Wyatt) with low (150mM) or high (1M) NaCl buffer conditions as described above. An aliquot of 100 µl sample of IMP2 at a concentration of 2 mg/ml +/− RNA was injected with a flow rate 0.5 ml/min. All experiments were performed at room temperature (25 °C). Data collection and SEC-MALS analysis were performed with ASTRA 8.0.0.19 (64-bit, Wyatt Technology). The refractive index of the solvent was defined as 1.3309 and the viscosity was defined as 0.8902. Dn/dc (refractive index increment) values for all samples was defined as 0.1850 (mL/g).

### Mass photometry (MP)

MP experiments were performed using a Reyfeyn Two MP Mass Photometer. Microscope coverslips and chambers were cleaned with isopropanol (HPLC grade) and Milli-Q H2O (5 min each), followed by drying and assembly. Apoferritin (ApoF), β-amylase (BAM), and Thyroglobulin (TG) were used as standards and were measured right before measurements of protein +/− RNA for both low (150mM) or high (1M) NaCl conditions. Protein + RNA conditions were incubated at room temperature for 1 hour before runs. Each protein was measured in new flow-chambers (i.e., each flow-chamber was used once). To acquire dynamic mass photometry movies, 15 µL of fresh buffer was first placed on the sample stage to optimize the focus of the microscope. The focal position was identified and locked in place with an autofocus system based on total internal reflection for the entire measurement. For each acquisition, 5 µL of protein was introduced into the flow-chamber (to achieve final working concentration of 20 nM) and measurements were recorded. Data acquisition was started right after (≤ 5s) the addition of protein using AcquireMP (Refeyn Ltd, v2.3). Images were time averaged 5-fold and pixel binned 4 × 4 before saving, resulting in an effective pixel size of 84.4 nm and effective frame rate of 200 Hz. DiscoverMP (Refeyn Ltd, v2.3) was used to generate the standard curves and data analysis.

### Size-exclusion chromatography coupled to small angle X-ray scattering (SEC-SAXS)

Summary of SAXS data collection and analysis tools can be found in **Table S3**.

#### SEC-SAXS data collection and processing

Samples were loaded into a 96-well auto-sampler plate prior to injection into a pre-equilibrated size exclusion-coupled small-angle X-ray scattering (SEC-SAXS) system at the center LiX Beamline of the NSLS-II located at Brookhaven National Laboratory (BNL)(39, 40). An Agilent 1260 Infinity Bio-Inert high-performance liquid chromatography (HPLC) system with an auto-sampler was used for sample injection and SEC. The beamline was configured to a 1.1 °A X-ray wavelength and 1.5mm path-length to obtain the relevant wave-vector, Q = 4sin(θ)/λ, where 2θ is the scattering angle and λ is the X-ray wavelength, yielding a q-range from 0.006 to 3.2 °A. A Superdex 200 increase 5/150 column (Cytiva) was equilibrated with buffers and used to separate proteinaceous species using a flow-rate of 0.35 mL/min during 2 s X-ray exposures over the course of 15-minute SAXS data collections. SAXS/WAXS images were radially integrated before being background subtracted using LIX specific pytho packages *py4xs* and *lixtools.* Subtracted profiles were exported and imported for analysis in BioXTAS RAW (RAW) (41).

#### SAXS data analysis and modeling

Background subtracted SAXS profiles were analyzed in RAW during primary analysis and data assessment. The structural models of IMP2 as predicted by AlphaFold2 or deduced from HDX were used for modeling in fragment form, maintaining the core domains intact. Conformational sampling, scoring of conformers with the SAXS data, and enumeration of any resultant multi-state models of GrsA was performed using FoxS (42).

IMP2 structures were prepared for input to FoxS using CHARMM-GUI’s PDB Reader and Manipulator. The independent domains of IMP2 were allowed diffuse mobility with respect to one another as tethered by flexible loops connecting rigid bodies. The conformers were scored against the SAXS data with FoXS before multi-state ensembles enumerated using MultiFoXS. *χ*2 estimates were verified against the predicted theoretical scattering as standard in the field using FoxS (43).

The dimer stacking interaction was predicted using the online served HDOCK (http://hdock.phys.hust.edu.cn/). Individual dimers (.pdb files) were loaded as both ligand and receptor to test for homo-oligomeric interactions and docking. The primary result with the best docking score (relative) and least RMSD is presented in the manuscript with a confidence score of > 80%. The docking is performed in a non-templated manner to avoid any biasness from user-inputs. The subsequent tetramers formed were analyzed using PyMol.

## Data availability

The mass spectrometry proteomics data have been deposited to the ProteomeXchange Consortium via the PRIDE(44) partner repository with the dataset identifier PXD049065.

SAXS data is available at SASBDB with draft reference IDs 5616 (apo/RNA-free) and 5617 (RNA-bound).

All other data is available upon request from.

## ACKNOWLEDGEMENTS

We thank the members of the Janiszewska and Kojetin laboratories for their critical reading of this manuscript and useful discussions; Dr. Brian MacTavish and Dr. HaJeung Park for their technical expertise. We would like to thank the beamline scientists at Lawrence-Berkeley National Laboratory for SAXS measurements. This work was supported by start-up funds from the Scripps Research Institute (M.J.) and NIH grants R01AG070719 (D.J.K.) and R01DK124870 (D.J.K.). The LiX beamline is part of the Center for Biomolecular Structure (CBMS), which is primarily supported by the National Institutes of Health, National Institute of General Medical Sciences (NIGMS) through a P30 Grant (P30GM133893), and by the DOE Office of Biological and Environmental Research (KP1605010). LiX also received additional support from NIH Grant S10 OD012331. As part of NSLS-II, a national user facility at Brookhaven National Laboratory, work performed at the CBMS is supported in part by the U.S. Department of Energy, Office of Science, Office of Basic Energy Sciences Program under contract number DE-SC0012704.

## AUTHOR CONTRIBUTIONS

S.Z. designed the study, conducted experiments, analyzed data, and wrote the manuscript draft. P.M.T, X.Y, and M.N.K.G performed and/or assisted with protein expression, biophysical assays, and data analysis. T.O. performed HDX and analyzed data. A.H.H. and S.F. helped with molecular modeling. R.N.R. performed SAXS data analysis. P.R.G., D.J.K., R.R. and M.J. designed experiments and supervised the study. All authors contributed to writing the manuscript.

## DECLARATION OF INTERESTS

M.J. is a member of a scientific advisory board at ResistanceBio.

## Notes

### Competing Interest Statement

All authors declare no competing interests. M.J. serves on scientific advisory board at ResistanceBio.

