## Supplemental Figures and Tables for "Structural insights into IMP2 dimerization and RNA binding"

### SUPPORTING INFORMATION

#### SUPPLEMENTAL FIGURE AND TABLE LEGENDS

**Figure S1. IMP2 purification. A**) SDS-PAGE gel of a His-Trap purification in high salt condition (1 M NaCl). **B**) SDS-PAGE gel at final stage of IMP2 purification. **C**) Circular dichroism of full-length purified IMP2 protein. **D**) Superdex-200 FPLC UV trace demonstrating multiple states of IMP2 during protein purification. IMP2 monomer approximated elution volume corresponding to 81ml peak. IMP2 dimer approximated elution volume corresponding to 72ml peak. IMP2 tetramer approximated elution volume corresponding to 48ml peak. **E**) Superdex-200 FPLC UV trace demonstrating IMP2 in mostly a dimeric state after protein purification; 72ml elution peak.

**Figure S2. CD thermal denaturation of IMP2 in low (150mM) and high (1M) salt conditions.** Apo/RNA-free IMP2 thermal melt. The data are also plotted in Fig. 5C and D.

**Figure S3. Tryptophan intrinsic fluorescence.** Fluorescence quenching assay, excitation at 295nm.

**Figure S4. Electrostatic surface maps of stacked dimers.** Two IMP2 dimers are shown. Colored scale represents the electrostatic potential.

**Figure S5: Electric mobility shift assay (EMSA). A**) IMP2 and Cox7b RNA, RNA-stained gel. **B**) IMP2 and Cox7b RNA, Coomassie stained gel. **C**) IMP2 and VSV RNA, RNA-stained gel. **D**) IMP2 and VSV RNA, Coomassie stained gel. Corresponding lanes: 1) 15uM RNA only, 2) 75uM IMP2 only, 3) 15μM RNA + 3.75μM IMP2, 4) 15μM RNA + 7.5μM IMP2, 5) 15μM RNA + 15μM IMP2, 6) 15μM RNA + 30μM IMP2, 7) 15μM RNA + 45μM IMP2, 8) 15μM RNA + 75μM IMP2.

**Table S1. SEC-MALS and MP results.**

**Table S2. Zeta potential of IMP2 at 150mM and 1M NaCl.**

**Table S3. SAXS summary.**

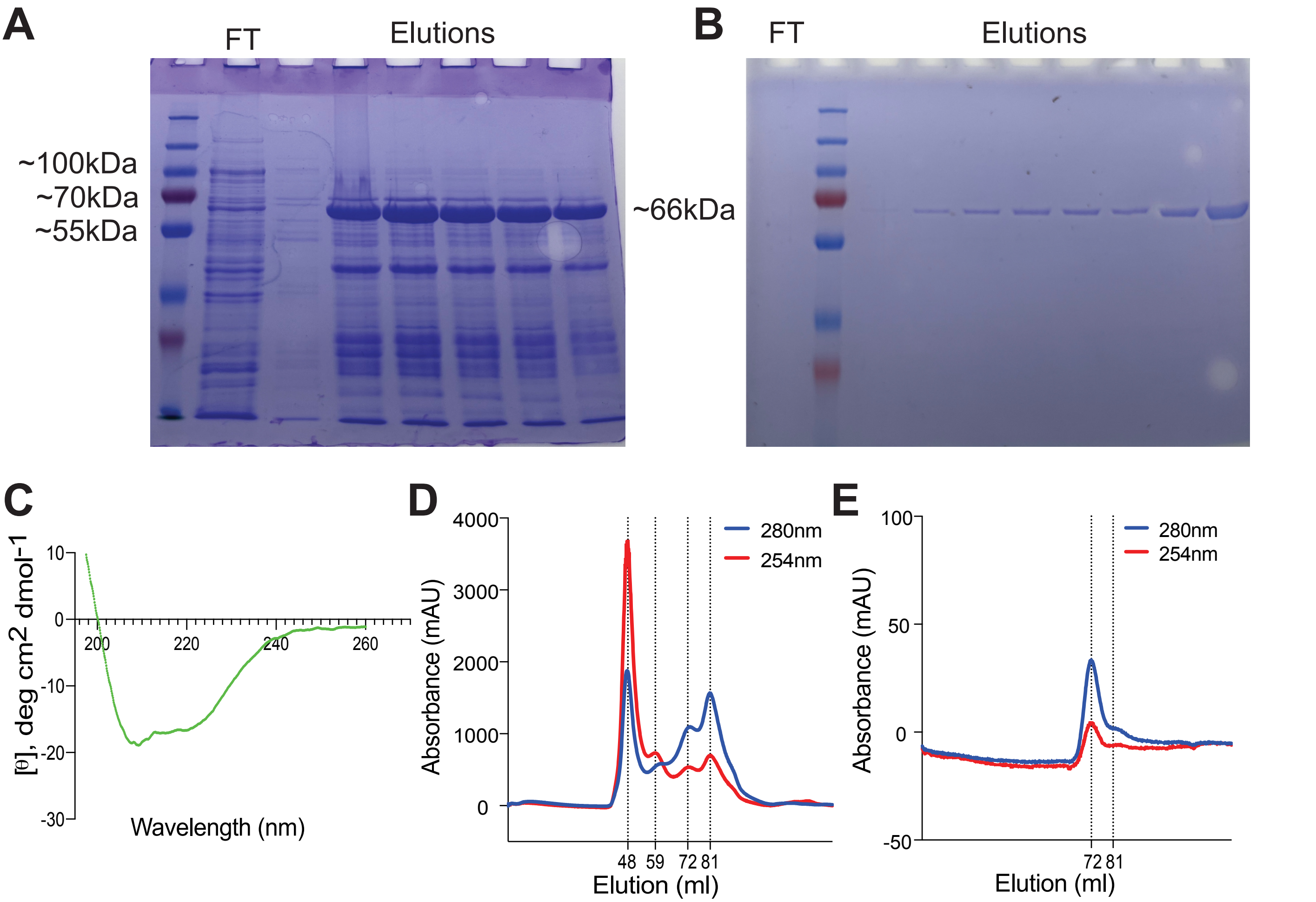
A
FT
Elutions
~100kDa
~70kDa
~55kDa
~66kDa
B
FT
Elutions
C
[θ], deg cm2 dmol-1
10
0
-10
-20
-30
200
220
240
260
Wavelength (nm)
D
Absorbance (mAU)
4000
3000
2000
1000
0
280nm
254nm
48
59
72
81
Elution (ml)
E
Absorbance (mAU)
100
50
0
-50
280nm
254nm
72
81
Elution (ml)

**Figure S1.**

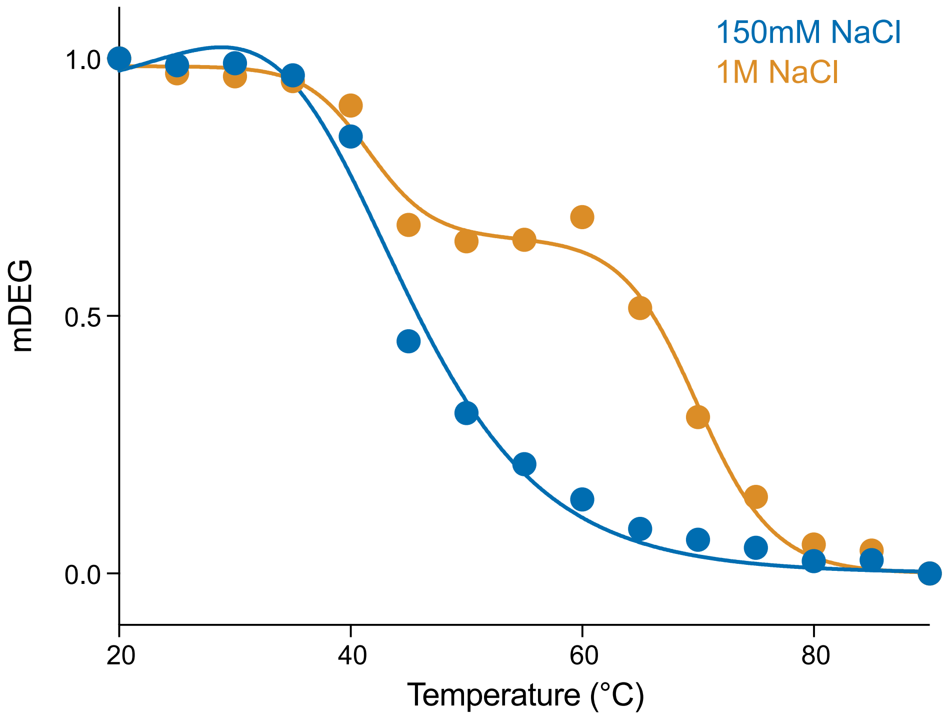


**Figure S2.**

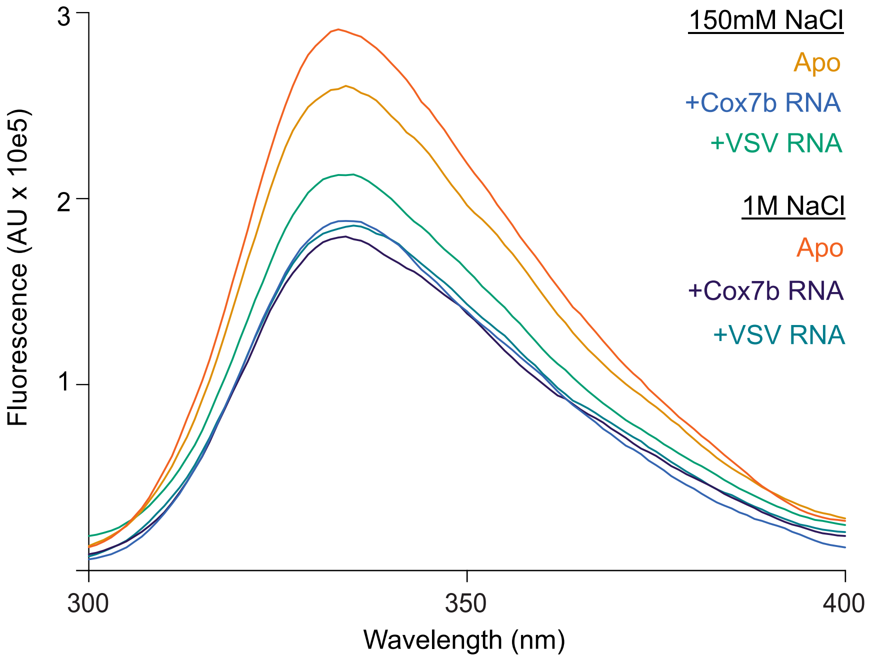

3
2
1
Fluorescence (AU x 10e5)
300
350
400
Wavelength (nm)
150mM NaCl
Apo
+Cox7b RNA
+VSV RNA
1M NaCl
Apo
+Cox7b RNA
+VSV RNA


### Figure S3.

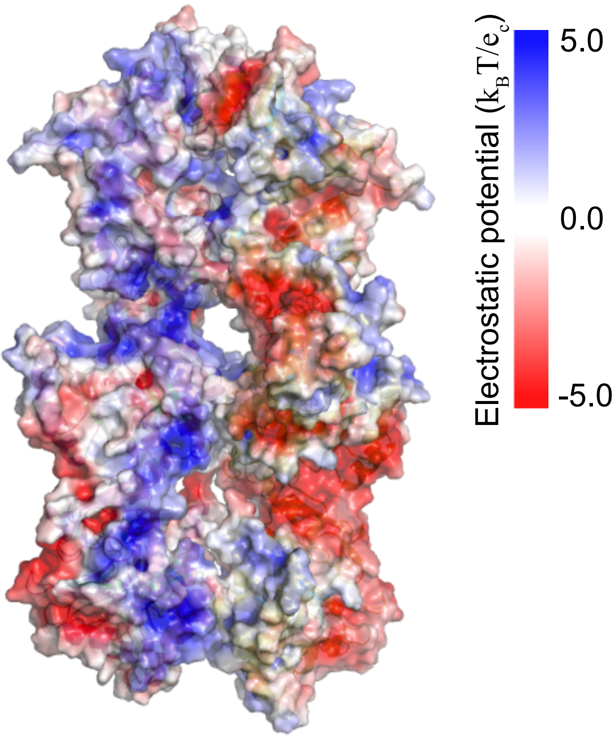


**Figure S4.**

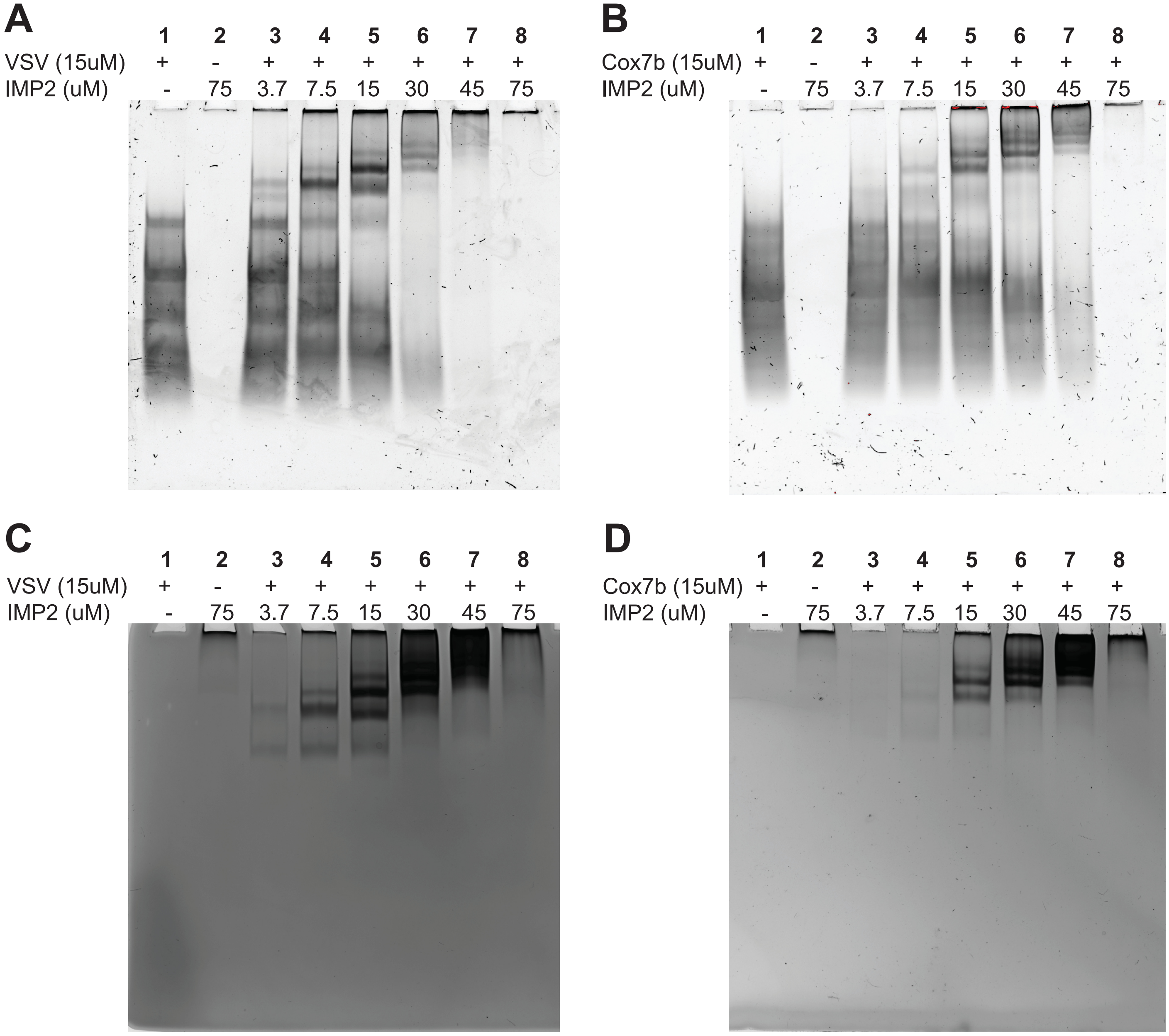
A
1 2 3 4 5 6 7 8
VSV (15uM) + - + + + + + +
IMP2 (uM) - 75 3.7 7.5 15 30 45 75
B
1 2 3 4 5 6 7 8
Cox7b (15uM) + - + + + + + +
IMP2 (uM) - 75 3.7 7.5 15 30 45 75
C
1 2 3 4 5 6 7 8
VSV (15uM) + - + + + + + +
IMP2 (uM) - 75 3.7 7.5 15 30 45 75
D
1 2 3 4 5 6 7 8
Cox7b (15uM) + - + + + + + +
IMP2 (uM) - 75 3.7 7.5 15 30 45 75

**Figure S5.**

| | | Mass Photometry | | | SEC-MALS | |
|---|---|---|---|---|---|---|
| **Buffer** | **Condition** | **Species 1** | **Species 2** | **Species 3** | **Species 1** | **Species 2** |
| 150mM NaCl | APO IMP2 | 127kDa (7%) σ 3.2kDa | 160kDa (82%) σ 9.2kDa | | 363kDa (±5.050%) | 785kDa (±6.388%) |
| | IMP2 + Cox7b RNA | 134kDa (77%) σ 4.7kDa | 160kDa (20%) σ 16.1kDa | | 406kDa (±0.091%) | |
| | IMP2 + VSV RNA | 131kDa (27%) σ 4.7kDa | 158kDa (54%) σ 13.3kDa | 233kDa (11%) σ 49kDa | 375kDa (±0.182%) | |
| 1mM NaCl | APO IMP2 | 120kDa (48%) σ 5.4kDa | 150kDa (42%) σ 12.8kDa | | 288kDa (±1.547%) | |
| | High: IMP2 + Cox7b RNA | | 143kDa (105%) σ 18.1kDa | | 85kDa (±0.819%) | |
| | High: IMP2 + VSV RNA | 131kDa (45%) σ 6kDa | 153kDa (43%) σ 11kDa | 254kDa (9%) σ 59kDa | 125kDa (±0.551%) | |

**Table S1.**

| | **IMP2 150mM NaCl** | **IMP2 1M NaCl** |
|---|---|---|
| | Mean | Mean |
| Zeta Potential (mV) | -2.658 | 5.077 |
| Conductivity (mS/cm) | 13.58 | 68.46 |
| Quality Factor | 0.129 | 0.3155 |

**Table S2.**

| Data Collection | Specifications |
|---|---|
| Beamline | LiX Beamline, NSLS-II |
| Wavelength of X-ray | 1.1 |
| Path Length (mm) | 1.5 |
| Detector | 3 x Pilatus 3 |
| Exposure Time (s) | 2 |
| q range ($Å^{-1}$) | 0.006 to 3.2 |
| Temperature (K) | 277.15 |
| **Size Exclusion Chromatography** | **Specifications** |
| Column | Cytiva Superdex increase 200 5/150 |
| Flow rate (ml/min) | 0.5 |
| Buffer composition (added substrates where needed) | 20mM Hepes pH 7.4, 150mM NaCl, 2mM MgCl2, 10% Glycerol |
| Temperature (K) | 277.15 |
| **Data Analysis Software** | |
| Primary data reduction | python package py4xs |
| Data processing | RAW |
| Rigid Body modelling | FoxS |
| 3D graphics representations | CHIMERA |

**Table S3.**
